# Integrated metabolic modeling, culturing and transcriptomics explains enhanced virulence of *V. cholerae* during co-infection with ETEC

**DOI:** 10.1101/2020.06.16.156158

**Authors:** Alyaa M. Abdel-Haleem, Vaishnavi Ravikumar, Boyang Ji, Katsuhiko Mineta, Xin Gao, Jens Nielsen, Takashi Gojobori, Ivan Mijakovic

**Affiliations:** King Abdullah University of Science and Technology (KAUST), Computational Bioscience Research Centre (CBRC), Thuwal, 23955-6900, Saudi Arabia; King Abdullah University of Science and Technology (KAUST), Biological and Environmental Sciences and Engineering (BESE) division, Thuwal, 23955-6900, Saudi Arabia; Novo Nordisk Foundation Center for Biosustainability, Technical University of Denmark, Kongens Lyngby, Denmark; Systems and Synthetic Biology, Department of Chemical and Biological Engineering, Chalmers University of Technology, Gothenburg, Sweden

**Keywords:** Infectious diseases, cholera, diarrhea, co-infection, drug target, flux balance analysis, constraint-based model, genome scale reconstruction

## Abstract

Gene essentiality is altered during polymicrobial infections. Nevertheless, most studies rely on single-species infections to assess pathogen gene essentiality. Here, we use genome-scale metabolic models to explore the effect of co-infection of the diarrheagenic pathogen *Vibrio cholerae* (*V. cholerae*) with another enteric pathogen, enterotoxigenic *E. coli* (ETEC). Model predictions showed that *V. cholerae* metabolic capabilities were increased due to ample cross-feeding opportunities enabled by ETEC. This is in line with increased severity of cholera symptoms known to occur in patients with dual-infections by the two pathogens. *In vitro* co-culture systems confirmed that *V. cholerae* growth is enhanced in co-cultures relative to single-cultures. Further, expression levels of several *V. cholerae* metabolic genes were significantly perturbed as shown by dual RNAseq analysis of its co-cultures with different ETEC strains. A decrease in ETEC growth was also observed, probably mediated by non-metabolic factors. Single gene essentiality analysis predicted conditionally-independent genes that are essential for the pathogen’s growth in both single- and co-infection scenarios. Our results reveal growth differences that are of relevance to drug targeting and efficiency in polymicrobial infections.

**Importance:** Most studies proposing new strategies to manage and treat infections have been largely focused on identifying druggable targets that can inhibit a pathogen’s growth when it is the single cause of infection. *In vivo*, however, infections can be caused by multiple species. This is important to take into account when attempting to develop or use current antibacterials since their efficacy can change significantly between single and co-infections. In this study, we used genome-scale metabolic models (GEMs) to interrogate the growth capabilities of *Vibrio cholerae* (*V. cholerae*) in single and co-infections with enterotoxigenic *E. coli* (ETEC), which co-occur in large fraction of diarrheagenic patients. Co-infection model predictions showed that *V. cholerae* growth capabilities are enhanced in presence of ETEC relative to *V. cholerae* single-infection, through cross-fed metabolites made available to *V. cholerae* by ETEC. *In vitro*, co-cultures of the two enteric pathogens further confirmed model predictions showing an increased growth of *V. cholerae* in co-culture relative to *V. cholerae* single-cultures while ETEC growth was suppressed. Dual RNAseq analysis of the co-cultures also confirmed that the transcriptome of *V. cholerae* is distinct during co-infection compared to single infection scenarios where processes related to metabolism were significantly perturbed. Further, *in silico* gene-knock out simulations uncovered discrepancies in gene essentiality for *V. cholerae* growth between single and co-infections. Integrative model-guided analysis thus identified druggable targets that would be critical for *V. cholerae* growth in both single and co-infections, thus designing inhibitors against those targets would provide a broader spectrum coverage against cholera infections.

## Introduction

Many studies focus on single-species infections although pathogens often cause infections as part of multi-species communities^1^. Most studies that aim at identifying essential genomes, for example, have largely depended on single cultures^2-5^. Such studies, thus, identify sets of ‘conditionally-dependent essential’ genes depending on the investigated growth conditions. Co-infecting microorganisms alter pathogen gene essentiality during polymicrobial infections^1^. Nevertheless, a limited number of studies have attempted to identify variations in growth capabilities or gene essentiality of a pathogen in co-infection conditions.

Many metabolic processes are critical for cellular growth and survival, and hence a pathogen’s anabolic and catabolic capabilities are usually tightly linked to its growth capabilities. There is growing evidence that, in addition to signals from the environment, the metabolism of a pathogen plays a major role in its virulence as well^6-9^.

Genome-scale metabolic network reconstructions^10-12^ (GEMs) have proven to be powerful tools to probe the metabolic capabilities of several enteric pathogens including *E. coli*^13^, *Shigella*^13^ and *Salmonella*^*14*^. GEMs are knowledge bases describing metabolic capabilities and the biochemical basis for entire organisms^10-12^. GENREs can be mathematically formalized and combined with numerical representations of biological constraints and objectives to create genome-scale metabolic models (GEMs)^10-12^. These GEMs can be used to predict biological outcomes (e.g. gene essentiality, growth rate) given an environmental context (e.g. metabolite availability^14,15^). Metabolic models recapitulate the biological processes of nutrient uptake and metabolite secretion, which can be the basis of some microbial interactions^16^. Growing number of experiments illustrated the predictive power of metabolic-driven computational approaches to describe emergent behaviors of co-existing species^17-22.^ However, deploying computational models to predict variations in pathogens’ growth capabilities when present in single or co-infecting scenarios has not been investigated.

*Vibrio cholerae* (*V. cholerae*) is a Gram-negative bacterium that causes acute voluminous diarrhea representing a dramatic example of an enteropathogenic invasion. Cholera infections are typically caused by contaminated food and water^23,24^. Seven cholera pandemics have been recorded in modern history and the latest is still ongoing^25,26,27^. *V. cholerae* life cycle is marked by repetitive transitions between aquatic environments and the host gastrointestinal tract, thus it has to adjust to different qualities and quantities of nutrient sources^28^. Within the human host, a highly active metabolic program is necessary to support *V. cholerae* high growth rates^28^ where it was reported that cell numbers reach up to 10^9^ cells/g stool excreted by cholera patients^23,28,29^. Further, several reports have suggested a role for central metabolism in regulating the production of virulence factors in *V. cholerae* (cholera toxin ‘CTX’, and toxin coregulated pilus ‘TCP’). For instance, TCP and CTX are not produced when *V. cholerae* is grown in M9-glycerol^30-32^. The Entner-Doudoroff pathway has been shown to be obligatory for gluconate utilization and plays an important role in regulating *V. cholerae* virulence^32^. While most case reports focus on *V. cholerae* as the single causative agent of diarrhea in case of Cholera infections, *V. cholerae* has commonly been involved in dual infections with enterotoxigenic *E. coli* (*ETEC*)^33-35^, the second most frequent cause (∼15%) of diarrheal diseases after *V. cholerae*. Notably, dual infections of *V. cholerae* and ETEC are associated with increased severity and increased healthcare costs^34^. Thus, there is a need to study the variations in growth capabilities and gene essentiality between single- and multi-species infections of pathogens in general, and of *V. cholerae* in particular.

Here, we built a *V. cholerae* genome-scale metabolic model and validated its single gene essentiality predictions against experimentally published data. We then evaluated the growth capabilities of *V. cholerae* in relation to other enteric pathogens by simulating their growth on 656 growth conditions spanning several nutrient sources under aerobic and anaerobic conditions. Following, we reconstructed a co-infection model of *V. cholerae* with ETEC in a shared environment and compared the growth capabilities of *V. cholerae* in single *vs*. co-infection settings. Co-infection model simulations allowed for a comprehensive assessment of variations in growth capabilities and single gene essentiality when *V. cholerae* is grown solely or in co-culture with ETEC. *In vitro* co-cultures of the two enteric pathogens as well as dual RNAseq data reflected corresponding variations in growth predictions and gene expression levels, respectively. Using single and co-infection models, we predicted *V. cholerae* essential genes representing potential druggable targets that would be of broader spectrum against *V. cholerae* both single and co-infections. The present work is computationally driven using high quality experimentally verified *in silico* and *in vitro* models, and can be viewed as a means to prioritize potential druggable targets of pathogens that are known to be involved in single and multi-species infections. Further, our results substantiate the notion that data-driven computational modelling coupled to experiments can predict and analyze microbial communities behavior.

## Results

### Characterizing the metabolic capabilities of *V. cholerae*

*i*AM-Vc960, a manually curated and quality-controlled GEM of *V. cholerae* was constructed (Figure 1, Step 1) to probe the enteric pathogen’s metabolic capabilities and gene essentiality in single and co-infections. We sequenced and annotated the genome of *V. cholerae* 52, an O37 serotype strain (see methods section, and Figure S1 at https://github.com/alyamahmoud/coinfection_modeling/blob/master/supplementary_material/supplementary_text.docx). A list of metabolic pathways in *V. cholerae* V52 was built based on the genome annotation generated in this study as well as those available in PATRIC and that of *V. cholerae* O1 N16961 (see Table S1 at https://github.com/alyamahmoud/coinfection_modeling/blob/master/supplementary_material/supplementary_tables.xlsx). The reconstruction was converted into a model and the stoichiometric matrix was constructed with mass and charge balanced reactions in the standard fashion using the COBRA toolbox v.3.0.^36^. Flux balance analysis was used to assess network characteristics and perform simulations^37^. The biomass function was constructed primarily based on that of *V. vulnificus*^7^ and *E. coli* K12 *i*JO1366^38^.Transcriptomics data of *V. cholerae* V52 single-cultures in minimal media was also generated and used to further refine *i*AM-Vc960 reconstruction and biomass objective function (see Table S1 at https://github.com/alyamahmoud/coinfection_modeling/blob/master/supplementary_material/supplementary_tables.xlsx). *i*AM-Vc960 accounts for 2172 reactions, 1741 metabolites across three compartments (cytosol, periplasm and extracellular compartments) and 960 metabolic genes. Gene-protein-reaction (GPR) associations could be defined for 72% of all enzymatic reactions (Figure 2A). *i*AM-Vc960 exceeds the automatically generated *V. cholerae* model as part of the Path2Models^39^ project in terms of its gene, metabolite and reaction content. 584 (89%) of the Path2Models *V. cholerae* model genes were already in *i*AM-Vc960. The remaining 68 genes were mostly non-metabolic. The Path2Models *V. cholerae* model as downloaded from the biomodels repository was unable to produce any biomass, thus we could not perform a functional comparison between *i*AM-Vc960 and the previously published *V. cholerae* model (see supplementary text at https://github.com/alyamahmoud/coinfection_modeling/blob/master/supplementary_material/supplementary_text.docx for details on comparison to other previously published *V. cholerae* GEMs^40^).

**Figure 1.**
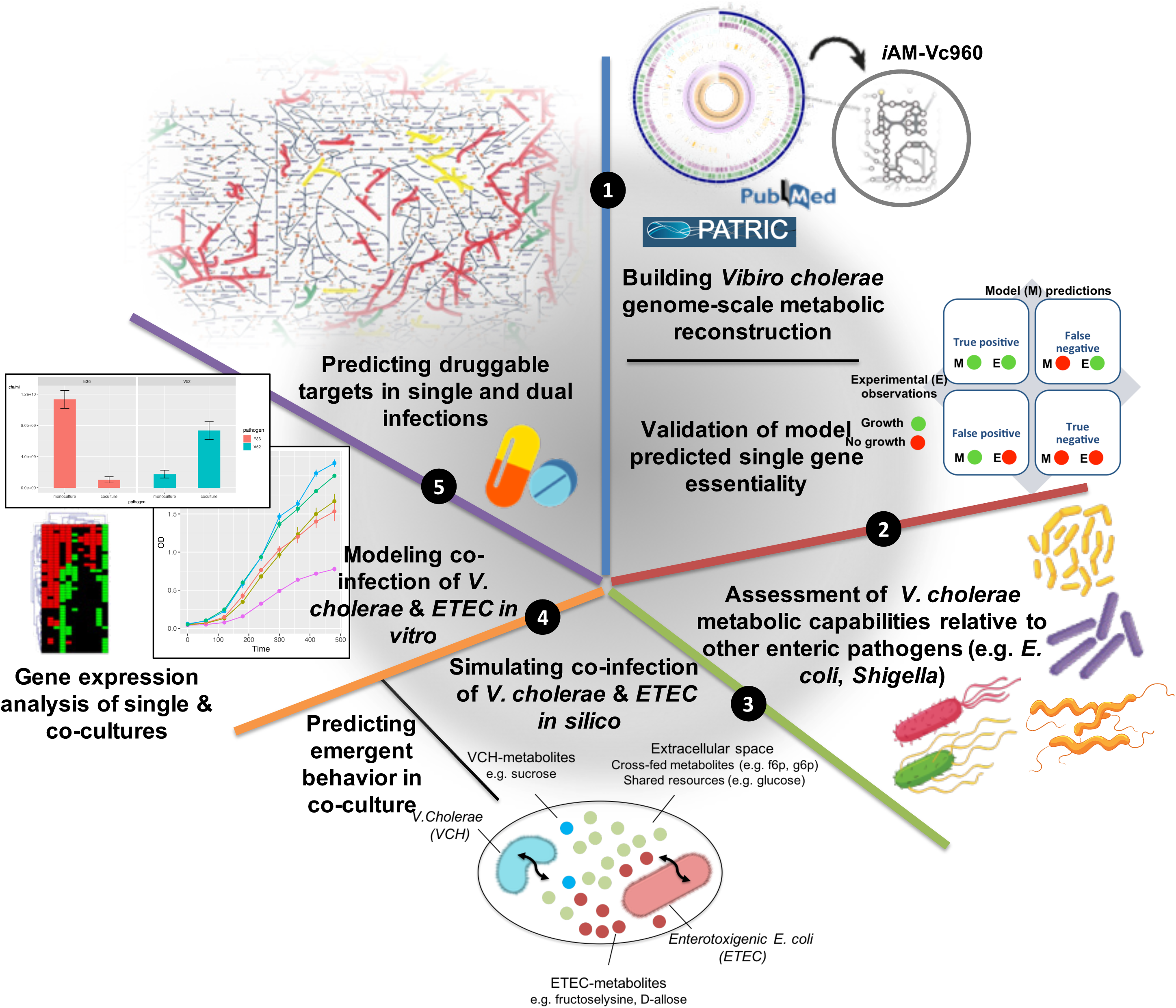
Overview of the study design.

**Figure 2.**
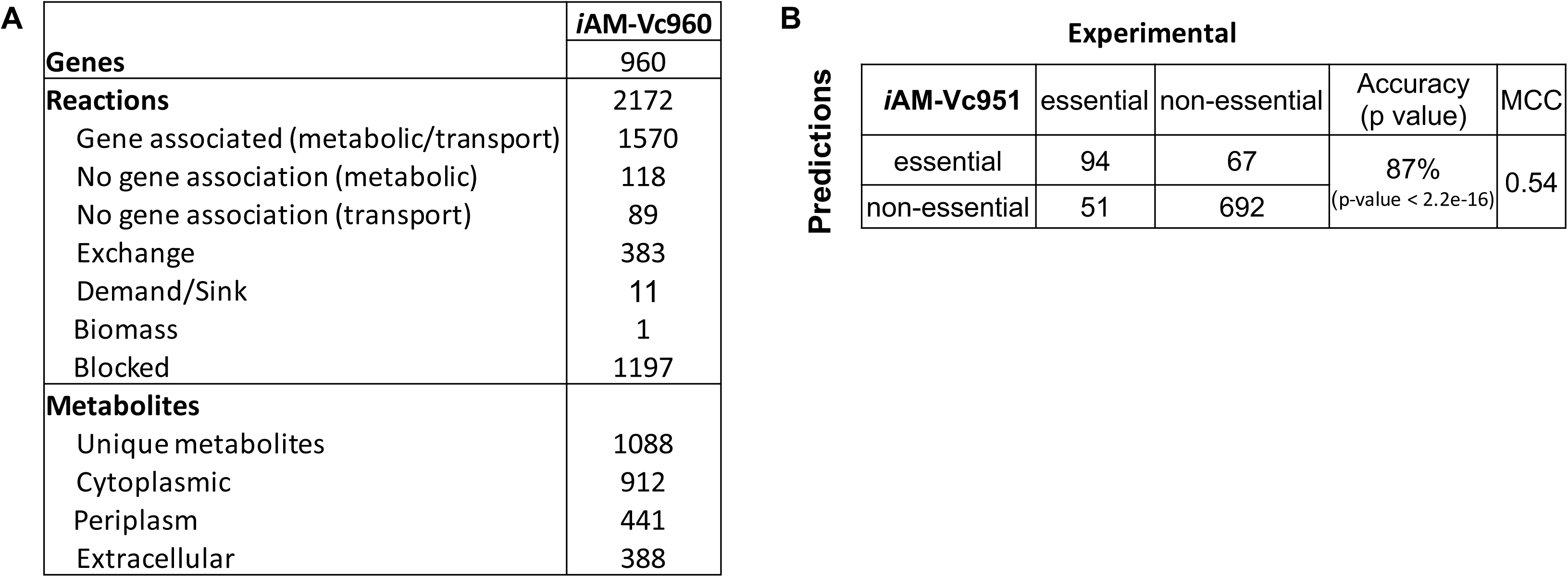
*V. cholerae* genome-scale model iAM-Vc960 description and performance evaluation. A) *V. cholerae* genome scale metabolic reconstruction stats (iAM-Vc959). C) Comparison of *i*AM-Vc960 gene essentiality predictions (simulating *in vitro* growth conditions in LB) showed 87% accuracy when compared to single gene deletion experiments from OGEE essential (n = 458) and non-essential (n = 758) gene datasets. *In silico* gene essentiality was graded according to the percentage of reduction in growth rate compared to wild type. Fisher exact test as well as Mathew correlation coefficient (MCC) were used to compute significance of overlapping consistent predictions for *i*AM-Vc959. See Table S2 at https://github.com/alyamahmoud/coinfection_modeling/blob/master/supplementary_material/supplementary_tables.xlsx for details.

*i*AM-Vc960 predicted growth rate was 1.07 mmol/gDW/h, in M9 minimal medium supplemented with glucose, corresponding to a doubling time of 39 minutes. Previous experiments^41^ using *V. cholerae* species reported doubling times of 38 min and 147 min for fast and slow growth, respectively. Hence, *i*AM-Vc960 predicted doubling time was within the expected range.

In order to further validate *i*AM-Vc960 predictions, we tested if *i*AM-Vc960 could correctly predict gene essentiality. Multiple attempts have been made to generate definitive lists of essential genes, but there are still many discrepancies between these studies even for a model bacterium such as *E. coli* strain K-12^42^. We thus compiled a high confidence set of genes (n = 223, see Table S2 at https://github.com/alyamahmoud/coinfection_modeling/blob/master/supplementary_material/supplementary_tables.xlsx) that have been shown to be critical for *V. cholerae* growth and survival from three independent previously published studies^43-45^. In rich medium (Luria-Bertani broth, LB), *i*AM-Vc960 correctly predicted 71% of the experimentally verified metabolic gene knock-outs (see Table S2 at https://github.com/alyamahmoud/coinfection_modeling/blob/master/supplementary_material/supplementary_tables.xlsx). In a second step, we also used gene essentiality data for *V. cholerae* str. C6706, a closely related O1 El Tor isolate, obtained from the Online GEne Essentiality (OGEE) database^4,5^ which contains information for essential (n = 458) and non-essential genes (n = 3144) (see supplementary text for a comment on serotype differences at https://github.com/alyamahmoud/coinfection_modeling/blob/master/supplementary_material/supplementary_text.docx). The overall accuracy of *i*AM-Vc960 in reproducing OGEE essentiality (and non-essentiality) data was 87% (Figure 2B) (see supplementary text at https://github.com/alyamahmoud/coinfection_modeling/blob/master/supplementary_material/supplementary_text.docx for details). Overall, *i*AM-Vc960 predicted 225 and 171 genes to be essential for optimal *V. cholerae* growth in minimal and rich media, respectively.

The agreement between the experimental gene essentiality data, obtained from previously published studies, and the computational results, generated in the current study, in terms of growth and single gene essentiality predictions, on the whole, validates the content of the reconstruction, the modeling procedure and the objective function definition (Figure 1, step 1). As such, *i*AM-Vc960 is a high-quality manually curated genome-scale model that can simulate *V. cholerae* metabolism and thus, can be used to predict phenotypic behavior of *V. cholerae* in response to different perturbations (e.g. culture conditions, interaction partners …etc). This prompted us to systematically and comprehensively assess the metabolic capabilities of *V. cholerae* to study how the pathogen adapts its network across the different growth conditions, assess the relative metabolic capacity of *V. cholera*e in relation to other enteric pathogens, as well as how the pathogen’s growth capabilities and gene essentiality is impacted in presence of other co-infecting pathogens.

### *V. cholerae* has restricted metabolic capabilities compared to *E. coli* and *Shigella*

Since enteric bacterial pathogens span several genera including *Escherichia, Salmonella*, and *Shigella*, we thought it would be relevant to assess the metabolic capabilities of *V. cholerae* in relation to other pathogens that cause diarrhea (Figure 1, Step 2). Using *i*AM-Vc960, we simulated growth capabilities of *V. cholerae* relative to a set of previously published^13^ GEMs of 55 *E. coli* (both commensal and pathogenic) and *Shigella* species on minimal media with 656 different growth-supporting carbon, nitrogen, phosphorous, and sulfur sources in aerobic and anaerobic conditions^13,14^. *i*AM-Vc960 model size was in line with the smaller genome size of *V. cholerae* compared to *E. coli* and *Shigella* (Figure 3A) where *V. cholerae* has 3855 ORFs while *Shigella* and *E. coli* each has on average 4199 and 4663 ORFs, respectively. Nevertheless, *i*AM-Vc960 metabolic genes covered 25% of *V. cholerae* ORFs^46^. Notably, *i*JO1366, the most well developed and curated genome-scale metabolic model covers 29% of *E. coli* str. K-12 substr. MG1655 ORFs. On average, *Shigella* and *E. coli* GEMs covered 27% and 29%, respectively of the corresponding species ORFs.

**Figure 3.**
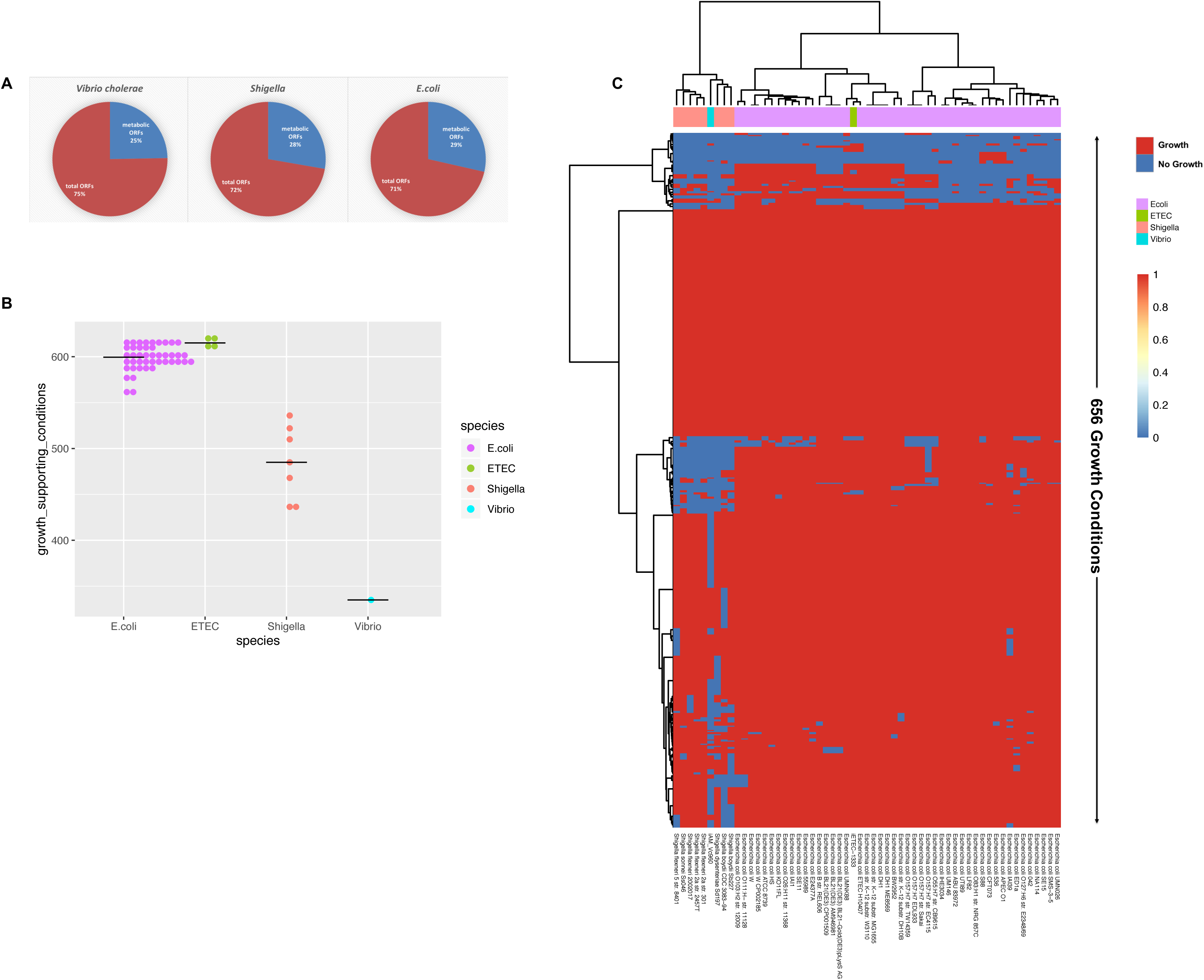
Functional assessment of *V. cholerae* metabolic capabilities relative to *E. coli* and Shigella. A) Proportion of metabolic genes included as GPR in GEMs of *E. coli* and Shigella^13^ and *V. cholerae* (this study) relative to total number of ORFs in each species. B-C) Assessment of *i*AM-Vc960 metabolic capabilities compared to a set of 55 *E. coli* and Shigella species^13^ by unique growth-supporting conditions. Predicted metabolic phenotypes on the variable growth-supporting nutrient conditions composed of different carbon, nitrogen, phosphorus, and sulfur nutrient sources in aerobic and anaerobic conditions. Strains were clustered based on their ability to sustain growth in each different environment. Columns in B represent individual strains, and rows represent different nutrient conditions. *i*AM-Vc960 co-clustered with *Shigella boydii* CDC 3083-94, *Shigella boydii* Sb227 and *Shigella dysenteriae* Sd197. Table S4-5 at https://github.com/alyamahmoud/coinfection_modeling/blob/master/supplementary_material/supplementary_tables.xlsx provide all details about the simulation conditions for the alternative nutrient sources. A growth rate of 0.01 was used as the cutoff for binarizing the simulation results and was used to construct the heatmap in B.

We first confirmed known metabolic differences for distinguishing *V. cholerae* from other enteric pathogens (Figure 3B-C). For instance, *i*AM-Vc960 predicted the ability of *V. cholerae* to utilize sucrose as sole carbon source^47,48^. *i*AM-Vc960 could not utilize arginine as sole carbon or nitrogen sources while all *E. coli* and *Shigella* models were able to utilize arginine under aerobic conditions^49,50^ in line with the frequent usage of the absence of arginine metabolism for characterizing *V. cholerae*^51^. Similarly, while *E. coli* and *Shigella* were able to utilize myo-inositol as sole phosphorus source, *i*AM-Vc960 predicted failure of *V. cholerae* to grow when no other phosphorus source is present in the medium^49^. Further, *i*AM-Vc960 also correctly predicted the ability of *V. cholerae* to utilize trehalose or mannitol as alternative carbon sources both in aerobic and anaerobic conditions^50,51^.

In contrast to *E. coli, V. cholerae* model displayed large loss of catabolic capabilities across the 656 tested growth conditions (Figure 3B-C, see Table S4-S5 at https://github.com/alyamahmoud/coinfection_modeling/blob/master/supplementary_material/supplementary_tables.xlsx). This computational result implies that *V. cholerae*, similar to *Shigella* and several pathogenic *E. coli*^52^, might have lost catabolic pathways for many nutrient sources. Model predictions showed that *V. cholerae* was able to grow in 51% (n = 336) of the simulated growth conditions, while *E. coli* and *Shigella* were able to grow, on average, in 92% (n = 602) and 75% (n = 493) of the tested growth conditions, respectively (Figure 3B, see Table S4-S5 at https://github.com/alyamahmoud/coinfection_modeling/blob/master/supplementary_material/supplementary_tables.xlsx) implicating that *V. cholerae* has less versatile metabolic capabilities compared to either *E. coli* or *Shigella*. In fact, *V. cholerae* metabolic capabilities were more similar to *Shigella* than *E. coli* (Figure 3C). *V. cholerae* model completely lost the capability to sustain growth on nutrient sources for which most of *E. coli* and *Shigella* models had growth capabilities. Some of these nutrients include D-lactate, D-fumarate, lactose, L-alanine-glutamate, uridine, xanthosine, thymidine, R-Glycerate, sn-Glycero-3-phosphoethanolamine, 4-Hydroxy-L-threonine, L-Asparagine, L-proline, L-Arabinose, and L-Xylulose as carbon sources as well as nitrate, nitrite^53^, ornithine, L-proline, agmatine, uracil, and putrescine^54^, as nitrogen sources, and myo-Inositol-hexakisphosphate as phosphorus sources. Further, most *Shigella* models and *i*AM-Vc960 were unable to sustain growth on chitobiose, D-Malate, D-Sorbitol, L-Fucose, ethanolamine, galactitol, propionate, D-Galactonate, choline, allantoin as sole carbon sources as well as hypoxanthine, inosine, and urea as nitrogen sources, whereas almost all other *E. coli* models examined were able to sustain growth under the same conditions.

Several tests based on nutrient utilization are routinely used to distinguish between pathogens that cause diarrhea. Using GEMs of enteric pathogens can aid in predicting potential metabolite markers that, upon experimental validation, could be used in clinical practice to diagnose the causative agent of diarrhea or an enteric pathogenesis in general.

### Predicted expanded growth capabilities of *V. cholerae* in co-culture with ETEC

Computational approaches modelling metabolic fluxes between organisms can be used to provide a mechanistic understanding of interaction patterns between different microbes^17,21,55,56^. An emergent behavior in co-culture will also relate to the extent of overlapping resources between the component species as well as whether or not there will be any cross-fed substrates^22^. Using *V. cholerae* as our model organism, we wanted to investigate how the metabolic capabilities (as proxy of growth capabilities) of *V. cholerae* will vary if other co-infecting pathogens are involved (Figure 1, Step 3). We thus set to model co-infections of *V. cholerae* and ETEC. *V. cholerae* (∼25%) followed by ETEC (∼15%) are the most prevalent bacterial pathogens causing diarrheal diseases in the developing world^33^. These bacteria are representative of species found in the same environment and are both involved in enteric pathogenesis. In particular, the choice of these species was inspired by the recurrent dual infections of both species in hospitalized patients due to diarrhea^33-35^. The antibody titer against cholera toxin (but not against heat-stable or heat -labile toxins produced by ETEC) was also found to increase in case of dual infections of *V. cholerae* and ETEC relative to single *V. cholerae* infections^34^, although no mechanistic explanation was attributed to these variations. *V. cholerae* V52 was also observed to be virulent against several other Gram-negative species including *E. coli* although ETEC was not tested^57^.

To investigate the behavior of the individual pathogens in co-infection relative to their single infections, we used *i*AM-Vc960 and a previously reconstructed GEM of ETEC, *i*ETEC1333^13^ to simulate the growth of *V. cholerae* and ETEC in a single shared environment^58,59^. Metabolic genes, metabolic reactions, and metabolites were compared across the species-specific networks. *i*AM-Vc960 and *i*ETEC-1333 had 1672 metabolites in common. This represented 96% and 85% of *V. cholerae* and ETEC total metabolites, respectively. To distinguish between shared and species-specific metabolites, each organism was represented as a separate compartment (Figure 4A) with a shared space representing the co-culture/infection medium. 23% (n = 380) of the common metabolites between the two models were amenable to exchange by being available in the shared extracellular space (Figure 4A). In total, the co-culture model, *i*Vc-ETEC-2293, had 4550 reactions, 3335 metabolites and 2293 genes. The objective function was set to maximize the biomass function of each pathogen, simulating growth of both species at 1:1 composition (see methods section and supplementary text at https://github.com/alyamahmoud/coinfection_modeling/blob/master/supplementary_material/supplementary_text.docx for details in development and refinement of the co-culture model).

**Figure 4.**
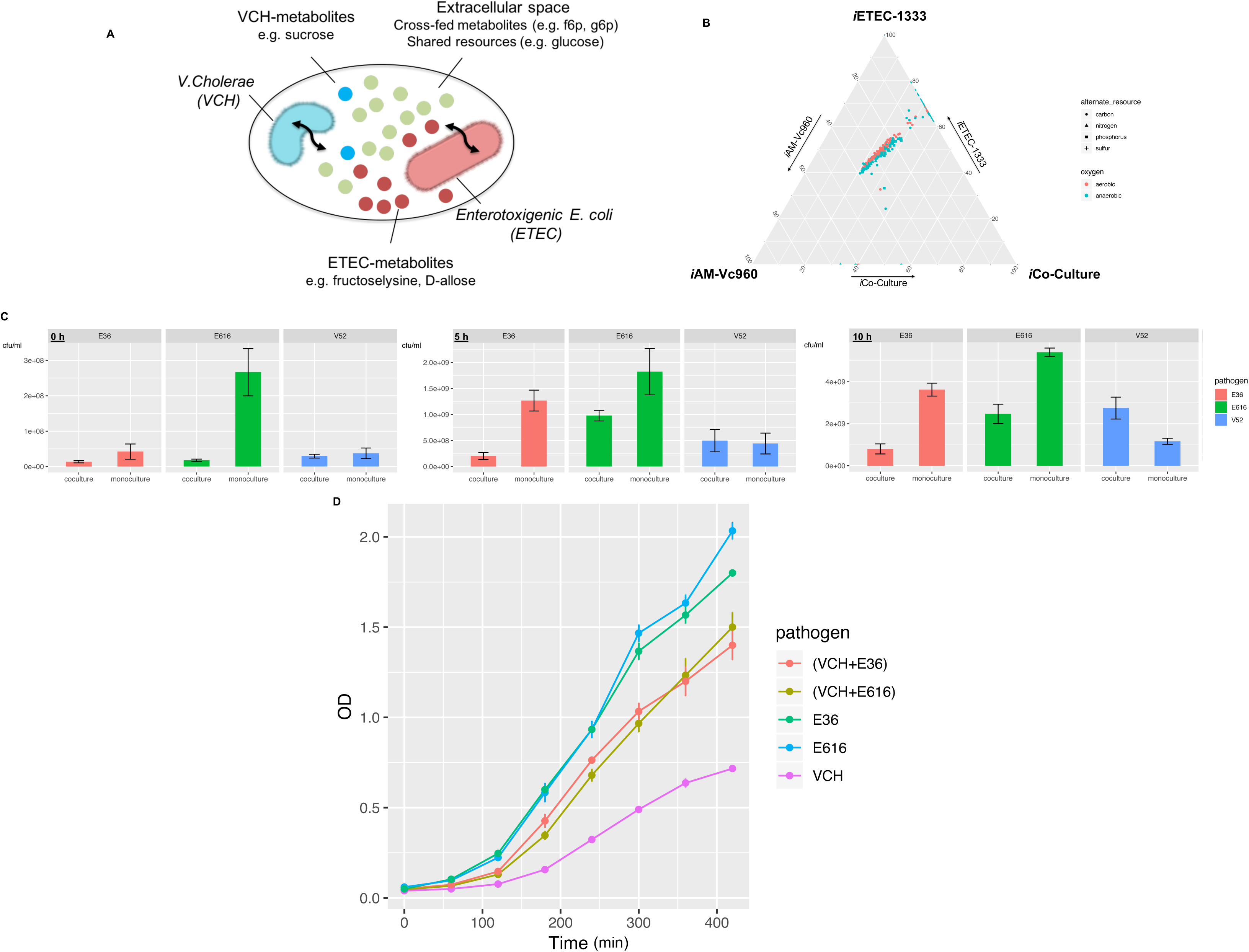
Computational modeling and *in vitro* co-culture of *V. cholerae* and ETEC co-infection. A) A Schematic showing the modeling framework used to simulate growth of *V. cholerae* and ETEC in a shared environment. B) Ternary plot showing +600 growth conditions to compare the metabolic capabilities of *V*.*cholerae* and ETEC monocultures relative to their co-culture. Values used for plotting are flux rates in the in the biomass objective function of each model and are meant to show the ability to grow or not grow under the respective growth condition rather than the flux value. No change in the overall plot was observed when using normalized values relative to standard growth conditions (aerobic conditions + glucose/ammonia/phosphate/sulfate. C) Quantification of *V. cholerae* and ETEC colony-forming units (CFUs) in monocultures and co-cultures over 10h for the CFU (pooled technical replicates of n = 3 biological replicates) in M9 minimal media supplemented with 0.5 % glucose, 1 mM MgSO4 and 0.1 mM CaCl2, 5uL spotted at each time point. Data shown as mean ± SD for three biological replicates. D) Dynamics of *V. cholerae* in co-culture with enterotoxigenic *E. coli* and in monoculture. Data shown as mean ± SD for three biological replicates

We then used the same set of 656 growth conditions to assess the difference in metabolic capabilities of *V. cholerae* and *ETEC* in single- and co-infections. All three models (*i*AM-Vc960, *i*ETEC1333 and *i*Vc-ETEC-2293) were able to grow in 51% (n = 333) of the tested growth conditions. ETEC was able to grow in 42% (n = 277) growth conditions that *V. cholerae* was unable to utilize in single-culture. However, *i*Co-Culture2993 acquired the capability to grow under the same conditions (Figure 4B, see Table S4-S5 at https://github.com/alyamahmoud/coinfection_modeling/blob/master/supplementary_material/supplementary_tables.xlsx). A closer look revealed that most of those acquired capabilities were due to ample cross-feeding opportunities enabled by the ETEC model. For instance, *i*AM-Vc960 is unable to grow on putrescine as sole nitrogen or carbon sources. *i*ETEC1333 and *i*Co-culture2993, however, are able to degrade putrescine into glutamate by putrescine transaminase (*patA*: ETEC_3343) or into glutamate and succinate through the gamma-glutamyl putrescine synthetase (*puuA*: ETEC_1401)/oxidoreductase (*puuB*: ETEC_1405) pathway, both being absent in *V. cholerae* genome. Similarly, *V. cholerae* cannot catabolize uridine (and xanthine) whereas ETEC can degrade uridine, xanthine and xanthosine into ribose as it possesses pyrimidine-specific ribonucleoside hydrolases (*RihA, RihB, RihC*: ETEC_0680, ETEC_2297, ETEC_0030) which can potentially be cross-fed to *V. cholerae*. In addition, several D-amino acids were observed to be cross-fed where they are degraded by ETEC into forms that can be utilised by *V. cholerae*, e.g. D-allose which is degraded by ETEC D-allose kinase (alsK: ETEC_4394) into fructose-6-phosphate that can be cross-fed to *V. cholerae*. Similarly, fructoselysine is metabolised by ETEC fructoselysine kinase (*frlD*: ETEC_3624) and fructoselysine 6-phosphate deglycase (*frlB*: ETEC_3622) into glucose-6-phosphate which can be cross-fed to *V. cholerae*. None of those genes have been identified in the genome of *V. cholerae* to date (determined via searching the annotated genome of *V. cholerae* O1 biovar El Tor str. N16961 in PATRIC^60^, Uniprot^61^, the annotated genome of *V. cholerae* V52 generated in this study as well as two other assemblies GCF_001857545.1 and GCF_000167935.2 retrieved through PATRIC^60^).

Overall, *i*ETEC1333, *i*AM-Vc960 and *i*Vc-ETEC-2293 were able to grow in 94% (n = 614), 51% (n = 336) and 93% (n = 613) of the simulated growth conditions, respectively (Figure 4B). As such, we predict that *V. cholerae* metabolic capabilities are expanded in co-infections with ETEC relative to *V. cholerae* single-infections while ETEC metabolic capabilities are almost not affected where the main differences among the two species lie in their capability to uptake and catabolize various nutrient sources. Our modeling approach thus provides mechanistic insights into the observed increase in cholera infection severity in clinical patients who demonstrated increased antibody titers against cholera (and not ETEC) toxin in case of co-infections by the two enteric pathogens^34^.

### Growth of *V. cholerae* is enhanced when co-cultured with ETEC *in vitro*

To validate our predictions, we employed single- and co-culture *in vitro* experiments (Figure 1, Step 4) to assess the predictions made by our enteric pathogens co-infection model (Figure 4C-D, see Table S6 at https://github.com/alyamahmoud/coinfection_modeling/blob/master/supplementary_material/supplementary_tables.xlsx). To this end, we developed a robust *in vitro* co-culture system of *V. cholerae* V52 and two different ETEC strains (E36 or E616) in M9 minimal medium supplemented with glucose (Figure 4C-D). All three tested strains (V52, E36 & E616) are clinical isolates that have been sequenced and characterized before^62,63^ (see supplementary text at https://github.com/alyamahmoud/coinfection_modeling/blob/master/supplementary_material/supplementary_text.docx for details on strain selection and sequencing performed as part of the current study). We determined the impact of the co-culture on each strain’s growth by comparing single culture abundance over 10 hours of growth to the abundance of each strain in co-culture at the same time (determined using CFU counting; all strains were in transition or stationary phase). E36 and E616 were shown to have diminished ability to grow in co-culture with *V. cholerae* V52. By contrast, growth of *V. cholerae* V52 was strongly enhanced in Co-culture conditions (Fig. 4C-D).

The growth data regarding *V. cholerae* V52 were in agreement with the modelling predictions. When comparing maximal abundances, cross-feeding and competitive interactions were already apparent. *V. cholerae* V52 reached higher maximal bacterial counts in *V. cholerae* V52/ETEC E36 (unpaired two-sided Wilcoxon: shift 5.8e+09, 90% confidence interval 3.8e+09 to 6.8e+09, p-value 0.07) and in *V. cholerae* V52/ETEC E616 (unpaired two-sided Wilcoxon: shift 5.6e+09, 90% confidence interval 4.4e+09 to 8.8e+09, p-value 0.1) co-cultures (Figure 4C-D). The maximum cell number of both ETEC strains tended to be lower when competing with *V. cholerae* V52 than when grown alone (unpaired two-sided Wilcoxon E36: shift -1.06e+10, 90% confidence interval -1.14e+10 to -8.60e+09, p-value 0.07; unpaired two-sided Wilcoxon E616: shift -6e+09, 90% confidence interval -9.4e+09 to -2.0e+09, p-value 0.1). Finally, according to maximal bacterial counts, E36 was more negatively affected by the presence of *V. cholerae* V52 than E616 (unpaired two-sided Wilcoxon E36: shift -6.4e+09, 90% confidence interval -9.4e+09 to -5.2e+09, p-value 0.1).

Although our modeling procedure predicted and explained the increase in *V. cholerae* growth capabilities when co-cultured with ETEC, the decrease in abundance of ETEC in *V. cholerae* V52/ETEC co-cultures was not captured by our metabolic models. *V. cholerae* V52 was previously found to be highly virulent against several Gram-negative bacteria, including *E. coli* and *Salmonella Typhimurium*, due to type VI secretion system (T6SS)^57^. Although ETEC was not tested for in these experiments, it is expected that ETEC would behave similarly to closely related pathogenic *E. coli* strains (EPEC, EHEC). Thus, the decrease in ETEC growth is very likely mediated by non-metabolic factors. We also focus on the improved growth of *V. cholerae* since this is of potential clinical relevance and since the decrease in ETEC growth in *V. cholerae* co-cultures has been investigated before.

### Altered gene expression in single- and multi-species co-cultures

To assess the level of genetic perturbations due to addition of ETEC as an interaction partner to *V. cholerae* cultures, we conducted a dual RNAseq analysis^64-67^ of *V. cholerae* co-cultures (Figure 1, Step 4) with each of the two ETEC strains (E36 or E616). We then compared the gene expression levels for each pathogen to its single-culture (see methods section and Tables S7-S10 at https://github.com/alyamahmoud/coinfection_modeling/blob/master/supplementary_material/supplementary_tables.xlsx). Through principal component analysis (PCA) (Figure 5, see Figure S5 at https://github.com/alyamahmoud/coinfection_modeling/blob/master/supplementary_material/supplementary_text.docx), we found that the co-cultures expression data clustered independently from single-culture data indicating that the transcriptome of *V. cholerae* is distinct during Co-culture compared to single-culture. The expression of 20% of *V. cholerae* quantifiable transcriptome was significantly altered when either strains of ETEC was added to the culture. In particular, 15-17% of *V. cholerae* genome was upregulated while 4-5% was downregulated in *V. cholerae* co-culture with ETEC relative to its single culture. *V. cholerae* differentially expressed genes were enriched in diverse metabolic processes spanning amino acid metabolism like tyrosine and L-phenylalanine (P value < 0.01, odds ratio > 10) as well as carbohydrate metabolic processes (P value < 0.05, odds ratio = 2.630409). (Figure 5, see Tables S9-S10 at https://github.com/alyamahmoud/coinfection_modeling/blob/master/supplementary_material/supplementary_text.docx). Upregulation of certain amino acid biosynthesis pathways, that can be catabolized by both species, highlights that despite potential cross-feeding between the two pathogens, presence of more than one infectious agent might eventually lead to competition^68^. Further, in support of non-metabolic mediated suppression in growth observed for ETEC, E36 differentially expressed processes were significantly enriched in taxis and chemotaxis GO terms (P value = 3.8e-05 and odds ratio > 20). Also, in line with previous reports^57,63^ about T6SS expression levels, T6SS components were constitutively expressed in *V. cholerae* V52 in both single- and co-cultures (see Tables S9-S10 at https://github.com/alyamahmoud/coinfection_modeling/blob/master/supplementary_material/supplementary_tables.xlsx).

**Figure 5.**
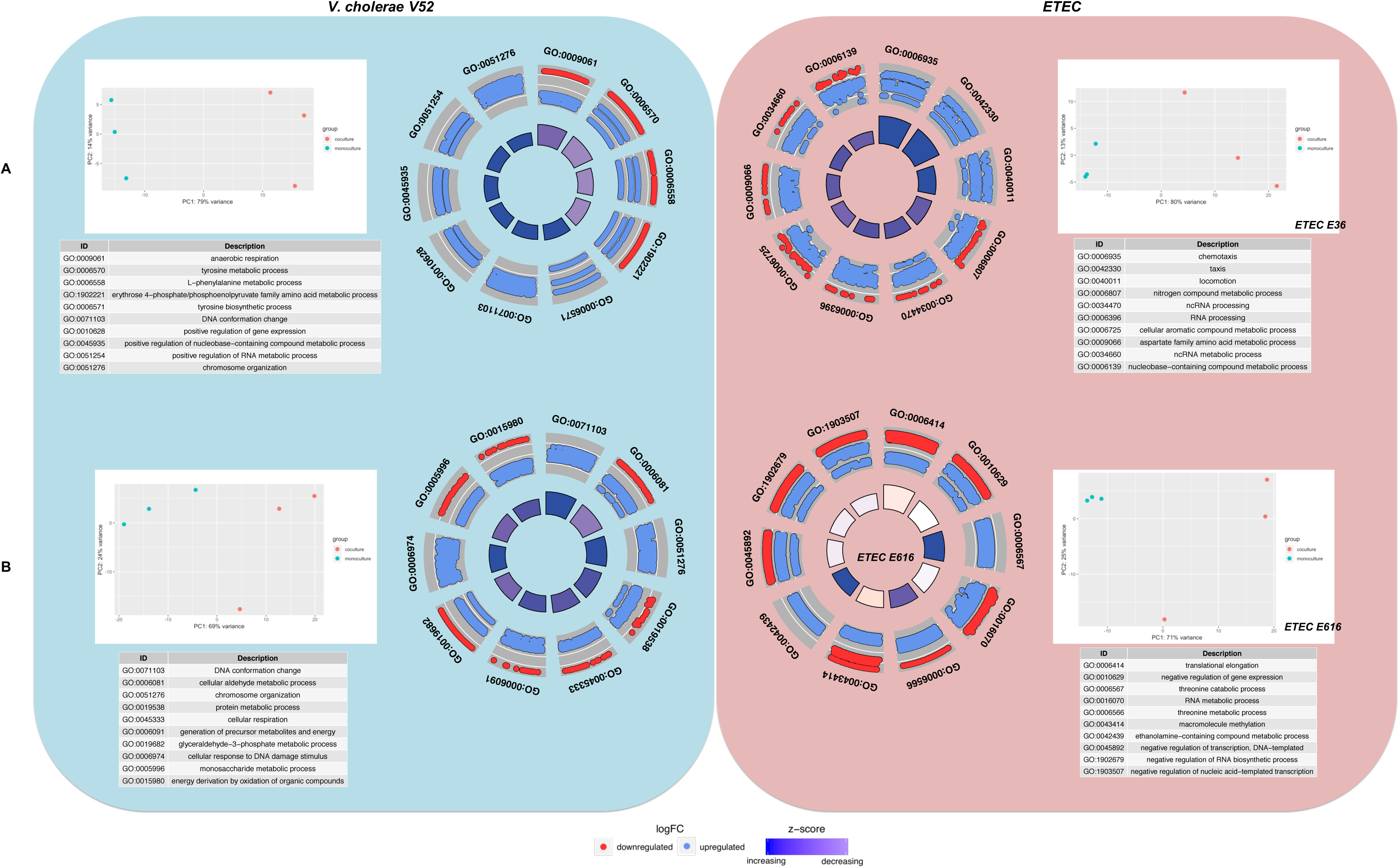
Dual RNAseq analysis of *V. cholerae* and ETEC in co-culture. GO enrichment of *V. cholerae* differentially expressed FIGfams in co-culture with ETEC A) E36 and B) E616 relative to its single culture. Up and down regulation is in *V. cholerae* when in co-culture relative to its mono-culture. Z-score is calculated according to GOplot (up-down/√*count*) where *up* and *down* are the number of assigned genes up-regulated (logFC>0) in the data or down-regulated (logFC<0), respectively.. PCA plots show that the mono-cultures are clustering differently from the co-cultures for either species.

In line with predicted cross-feeding interactions between *V. cholerae* and ETEC, we found that gamma-glutamyl putrescine oxidase (puuB), putrescine utilisation regulator (*puuR*) as well as several putrescine transporters were indeed significantly upregulated in E616/V52 co-culture relative to E616 single culture (logFC > 1.5, adjusted p value < 0.05). Furthermore, neither *patA* nor *puuB* were expressed in *V. cholerae* V52. Similarly, ribose 5-phosphate isomerase B (*rpiB*) and transcriptional regulator of D-allose utilization (*rpiR*) were significantly upregulated in E616/V52 co-culture relative to E616 single culture (logFC > 2, adjusted p value < 0.005) and were not expressed in *V. cholerae* V52. Lastly, transcriptional regulator of fructoselysine utilization operon (*frlR*), fructoselysine 6-kinase (*frlD*), fructoselysine 3-epimerase (*frlC*), and fructoselysine-6-phosphate deglycase (*frlB*) were also significantly upregulated in E616/V52 Co-culture relative to E616 single-culture (logFC > 1-1.5, adjusted p value < 0.05).

Interestingly, expression levels of bacteriocins’ related genes in ETEC strains showed that colicins’ production and tolerance genes were significantly upregulated in E616 co-culture with *V. cholerae* V52 relative to the individually grown E616 (see Table S8 at https://github.com/alyamahmoud/coinfection_modeling/blob/master/supplementary_material/supplementary_tables.xlsx). In contrast, E36, whose growth is more sensitive to co-growth with *V. cholerae* V52 failed to up-regulate genes encoding colicin V production and tolerance genes (see Table S7 at https://github.com/alyamahmoud/coinfection_modeling/blob/master/supplementary_material/supplementary_tables.xlsx). Colicin V is a peptide antibiotic that members of *Enterobacteriaceae* commonly used to kill closely related bacteria in an attempt to reduce competition for essential nutrients^69^. To sum up, the difference in expression level of genes encoding colicins production and resistance explains why E36 growth was more severely affected when co-cultured with *V. cholerae V52* relative to E616 (see Figure S5 at https://github.com/alyamahmoud/coinfection_modeling/blob/master/supplementary_material/supplementary_text.docx).

RNAseq thus confirmed that there is an emergent behavior in the co-cultures and that the observed changes were not just due to variations in inoculum composition or the lag phase^67^. Taken together, our integrated modeling, co-culturing and transcriptomics approach provided mechanistic insights into the observed increase in cholera infection severity in dual infections with ETEC where ETEC co-infection results in an increased growth of *V. cholerae* due to expanded metabolic capabilities enabled by ETEC. In parallel, *V. cholerae* suppress ETEC growth by non-metabolic factors resulting in an increase in cholera infection severity but not ETEC as monitored by antibody titer against species-specific toxins^34^.

### Evaluation of experimentally validated essential genes across single and co-infections models of *V. cholerae*

The essential genome of a large class of bacterial species has been characterized as it encodes potential targets for antibacterial drug development^42,70^. Interestingly, metabolic genes have predominated in studies of essential genomes of microbial pathogens^70,71^. With this in mind, we attempted to construct a comprehensive map of *V. cholerae* essential metabolic genome (Figure 1, Step 5) by projecting the list of experimentally validated essential genes onto our single and co-infection models’ predictions (Figure 6, see Table S2 https://github.com/alyamahmoud/coinfection_modeling/blob/master/supplementary_material/supplementary_tables.xlsx). Selecting targets that are critical in both single and co-infection settings would promote the discovery of novel targets or new combinations of existing antibacterials that would be effective in a broader spectrum of cholera infections. The color scheme of highlighted reactions (Figure 6) denotes model prediction classification across single and co-infections. The red group in Figure 6 highlights reactions predicted to be sensitive in both single as well as Co-infections; this is of particular importance since the efficacy of some of the commonly used treatment drugs might significantly be altered in presence of more than one infecting agent. There are several gene deletions associated with reactions for which drugs have not been developed (see Table S2 at https://github.com/alyamahmoud/coinfection_modeling/blob/master/supplementary_material/supplementary_tables.xlsx). These highlight potential targets for new drug development that may aid in treating enteric pathogenesis. We also note that the green group identifies reactions that were missed by the models, and highlights areas for future model refinement.

**Figure 6.**
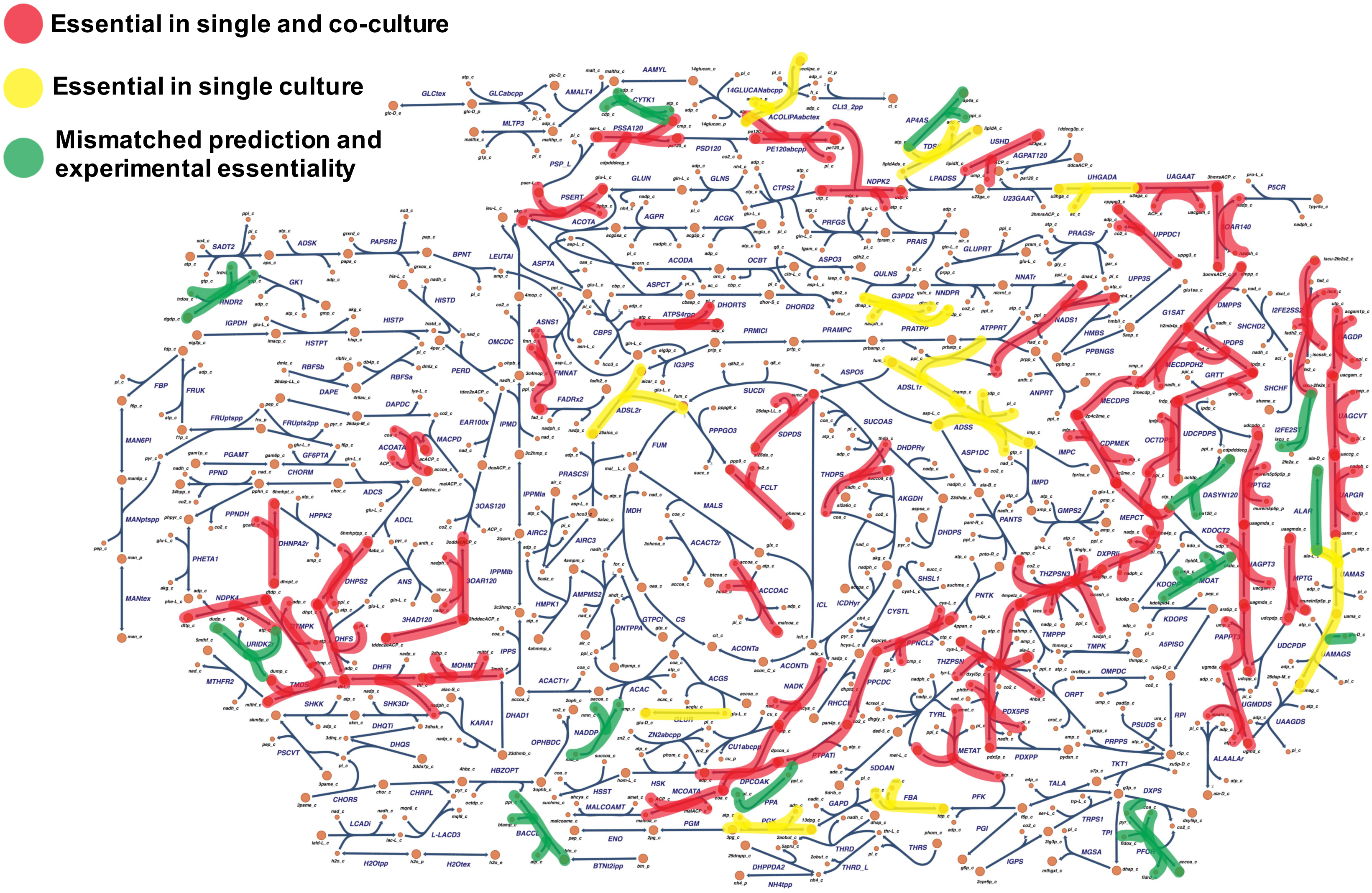
Comprehensive map of *V. cholerae* essential metabolic genome. constructed by projecting the list of experimentally validated essential genes onto our single and co-infection models’ predictions. Inhibitors against red-colored targets are expected to be of broader spectrum since they are be essential for *V. cholerae* in both single and co-infection scenarios. Inhibitors against yellow-colored targets are essential for V. cholerae growth in single-infection scenarios only losing their essentiality in presence of other co-infecting species. Green-colored targets are indicate a mismatch between model-predicted and experimentally-validated essentiality.

Out of the 80 metabolic genes that have been experimentally shown to be essential for *V. cholerae* growth and survival across several studies (see Table S2 at https://github.com/alyamahmoud/coinfection_modeling/blob/master/supplementary_material/supplementary_tables.xlsx), our co-culture model predicted 47 genes to be critical for *V. cholerae* growth even when a more metabolically versatile enteropathogen like ETEC is added to the culture irrespective of the variation in species composition (see methods section and Table S2 at for details). This set of 47 genes (Figure 6, red colored) represent potential drug targets that are predicted to be effective in killing *V. cholerae* whether it is the sole cause of diarrhea or as part of polymicrobial infection. Most of these enzymes were involved in cofactor biosynthesis (e.g. coenzyme A, tetrahydrofolate, FAD, pyridoxone-5-phosphate, pantothenate, and iron-sulfur cluster), isoprenoid and porphyrin metabolism as well as pyrimidine metabolism (Figure 6). Inhibitors of several of those enzymes have already been reported to have bactericidal effect^60^ in *V. cholerae* as well as in other enteric and non-enteric pathogens (see Table S2 at https://github.com/alyamahmoud/coinfection_modeling/blob/master/supplementary_material/supplementary_tables.xlsx). For instance, phosphopantetheine adenylyltransferase and thymidylate synthase have been already reported as drug targets^60^ in *V. cholerae* and ETEC E616. N-acetylglucosamine transferase is a promising drug target for *Salmonella enterica* while dephospho-CoA kinase has been shown to be an interesting drug target in ETEC E616 and *Shigella flexneri*^*60*^. Interestingly, 12 *V. cholerae* genes were also predicted to totally lose their essentiality in dual infections with ETEC. Some of those were involved in *de novo* purine metabolism (VC1126: *purB*, VC2602: *purA*) and carbohydrate degradation (VC0477: *pgk*, VC0478: *fbaA*) implicating that *V. cholerae* is probably depending on ETEC to salvage these nutrients.

ATP synthase subunits were essential for *V. cholerae* growth in single cultures as predicted by *i*AM-Vc960. Deletion of any of the 7 genes of F0/F1 ATP synthase locus in the co-culture model resulted in a species-composition dependent reduction in reduced optimal growth (see Table S2 at https://github.com/alyamahmoud/coinfection_modeling/blob/master/supplementary_material/supplementary_tables.xlsx). In models simulating high *V. cholerae* abundance relative to ETEC, ATP synthase subunits were essential for optimal co-culture growth. In contrast, models simulating higher ETEC abundance relative to *V. cholerae* were less affected when ATP synthase subunits were deleted. F0/F1 ATP synthase genes have been shown to be essential in a variety of bacteria^43,72-75^ and have been recently reported as essential in *V. cholerae*^43^. In *E. coli*, however, ATP synthase is not essential^43,76,77^. Thus, drug inhibitors (acting on ATP synthase subunits) that would normally kill *V. cholerae* in single-infections would have decreased efficacy in case of dual infections with *E. coli*. This suggests that comparison of essential genes between organisms can uncover distinct ecological and physiological requirements for each species^43^ and should inspire future experiments to validate our computational predictions. Similarly, sodium-dependent NADH dehydrogenase (Na+-NQR), a key component of the respiratory chain of diverse bacterial species, including pathogenic bacteria as well as succinate dehydrogenase subunits were also predicted to lose essentiality for *V. cholerae* growth when co-cultured with *E. coli*. Taken together, our *in silico* predictions of variations in essentiality between single and co-culture settings highlight the importance of considering both scenarios when prioritising druggable targets for downstream validation.

## Discussion

Using integrated metabolic modeling, *in vitro* culturing and transcriptomics, we investigated the growth phenotypes and single gene essentiality variations of a representative human pathogen, *V. cholerae*, when implicated in single or co-infections. We found that *V. cholerae* growth is enhanced in co-infection scenarios with ETEC. Our modeling procedures explained this increase in *V. cholerae* growth by an expansion in its metabolic capabilities through cross-fed metabolites enabled by ETEC, reproducing observed behavior in patients with dual infections by the two enteric pathogens. We further predicted a core set of essential genes that are critical for *V. cholerae* growth whether it is implicated in single or dual infections with ETEC.

Our modeling approach allowed us to chart possible metabolites that can be cross-fed to *V. cholerae* through ETEC (see Table S5 at https://github.com/alyamahmoud/coinfection_modeling/blob/master/supplementary_material/supplementary_tables.xlsx). Cross-feeding, in which one species produces metabolites consumed by another, has been shown more than often to be adopted by co-existing species across diverse environments^17,18,56,78^. Questions like: whether the release of cross-fed metabolites or byproducts would enhance or enable the growth of other species or whether it will be costless or associated with reduced fitness of the producer, are not usually clear. Such questions become of even greater importance when it comes to pathogens since this will have direct impact on the dosage and spectrum of used antibiotics. Our integrative approach provides insights into how to arrive at primary answers to similar questions that should direct future experimental work.

A large fraction of *V. cholerae* essential genome (36%) (see supplementary text at https://github.com/alyamahmoud/coinfection_modeling/blob/master/supplementary_material/supplementary_text.docx for details) consists of metabolic functions spanning several processes including cell wall biosynthesis, lipid metabolism, and cofactor biosynthesis^25-27^. Most essential genes for *V. cholerae* growth whether it is causing single-or co-infections were also involved in cofactor biosynthesis. Interestingly, cofactor-use-efficient pathways were often favored by organisms that depend on simple carbon sources under anaerobic conditions^79^ resembling growth conditions in the intestine^53,80^ where *V. cholerae* and ETEC establish their infection. The application of this work is of immediate relevance for the choice of antibiotics used in case of single-or polymicrobial infections. Strategies that depend on an increase in dosage of one drug or combining drugs of known efficacy against individual species might not necessarily work when two or more pathogens are operating together. Our findings indicate that the essential transcriptome of *V. cholerae* is distinct during co-infection compared to single-infection and highlight the importance of studying pathogen gene essentiality in polymicrobial infections. While replacement fluids are the main treatment line for cholerae infections, antibiotics are frequently used to lessen the diarrheal purging, decrease the need for rehydration fluids and shorten the recovery time^23^. For other human pathogens however, antibiotics are the main stay and we envision that our framework can be applied to other pathogens and their most frequently reported co-infecting partners. We believe that such an integrative approach could be routinely integrated as part of drug target development pipelines.

An integral part of constraint-based modeling relies on reconciling differences that arise between modeling and experiments^10-12,81^. In our case, co-infection models’ simulations predicted an increase in *V. cholerae* growth rate coupled to almost no-impact on ETEC growth capabilities. This is in line with recent studies showing that most organisms secrete a broad distribution of metabolically useful compounds without cost in a variety of environmental conditions^56^. However, our *in vitro* co-culture experiments revealed a significant decrease in ETEC growth rate leading us to conclude, in light of existing literature^57^, that the suppression in ETEC growth is potentially mediated by non-metabolic factors that are not captured by our GEMs.

Although our approach is based on computational predictions and *in vitro* experiments which definitely do not fully recapitulate *in vivo* conditions, our growth phenotype, predicted by Co-culture models and *in vitro* co-cultures, matched observed behavior in patients presenting with diarrhea while being co-infected with both *V. cholerae* and ETEC showing higher antibody titers against cholera toxin relative to patients infected with *V. cholerae* only^34^. Nevertheless, we realize that there are other processes that are not accounted for even after integrating data from various sources within the current approach. For instance, the fact that our metabolic model could not predict the decrease in ETEC growth rate implies that this effect is probably mediated by a non-metabolic factor that is not captured by the metabolic models as such. Future models, building upon the present reconstruction, can expand the modeling scope to account for synthesis and secretion of *V. cholerae* virulence factors^6-9^ in an attempt to investigate how the metabolic network of *V. cholerae* impacts the synthesis of its virulence factors. Co-culture experiments create an artificial community in a controlled environment and thus provide ideal conditions to test ecological concepts concerning community stability and dynamics that cannot easily be measured in macro-ecological complex systems^82^. However, most parts of the human intestine are hypoxic, vary in pH level^53,80^, and are inhabited by diverse sets of commensal microbes which are not accounted for when solely depending on *in vitro* experiments. Current predictions and experiments thus do not capture several of these factors including temperature, pH changes, signaling, gene regulation, serotype differences, and co-existing commensal microbes (which may account for the absence of the *V. cholerae* growth phenotype when using solid agar or spent media for co-infection modeling, see Figures S5-S6 and supplementary text at https://github.com/alyamahmoud/coinfection_modeling/blob/master/supplementary_material/supplementary_text.docx or details). Our study investigates a synthetic enteric pathogens community with a combination of *in vitro* single- and co-cultures, mechanistic modeling and gene expression analysis. Constraint-based modeling approaches, which can take emergent metabolism into account^37^, require high-quality metabolic reconstructions for each community member, which take months of curation effort to obtain^83^. However, the modular nature of the modeling approach followed here implies that such approaches can be scaled up to simulate polymicrobial infections as well as co-existing commensal microbes to further prioritize druggable targets that would be effective in even broader range of infection conditions and complex ecosystems. Collectively, this work illustrates the importance of harnessing the power of integrative predictive modeling coupled with Co-culture experiments to recognize potential amplification in a pathogen’s growth capabilities a priori which could contribute to downstream therapeutic and management options.

## Methods

The methods employed for the reconstruction, simulation, and analyses presented in this manuscript are briefly summarized below, with further details regarding the procedures, protocols, calculations, and quality control measures provided in the supplementary material. All supplementary tables are available as part of a github repository at : https://github.com/alyamahmoud/coinfection_modeling.

### Growth assays and CFU measurements

Bacterial strains were grown in M9 (Sigma Aldrich) minimal medium supplemented with 0.5 % glucose, 1 mM magnesium sulphate and 0.1 mM calcium chloride, unless otherwise specified. *V. cholerae* V52 (O37 serogroup) and the enterotoxigenic *Escherichia coli* strains (ETEC E616 and ETEC E36) were a kind gift from Prof. Sun Nyunt Wai, Umeå University, Sweden. *V. cholerae* and ETEC were grown either individually (mono-cultures – V52, E616 and E36) or in combination (co-cultures – V52/E616 and V52/E36) at 37 °C at 200 rpm. Co-cultures were started with equal concentrations of each strain. The absorbance (OD_600_) was measured every 1h over a period of 7h for the growth curve measurements. Simultaneously, at every hour, an aliquot was taken from each culture flask, serially diluted and 5µL were spotted (three technical replicates) on agar plates containing appropriate antibiotics (100µg/mL of rifampicin or 15µg/mL of tetracycline).V52 mono-cultures were spotted on rifampicin plates whereas ETEC E616 and E36 mono-cultures were spotted on tetracycline plates. Following, all co-cultures were spotted on both sets of antibiotic plates to distinguish between the individual strains during co-cultures. All plates were incubated for a period of 12-16h at 37 °C after which the colonies were counted and the CFU/mL value was calculated.

### DNA extraction, sequencing and genome assembly

Genomic DNA and plasmids (in case of ETEC) were extracted from bacterial cells for the purpose of whole genome sequencing. *V. cholerae* and ETEC cells (mono-cultures) were inoculated in rich LB (Sigma Aldrich) medium and grown at 37 °C at 200 rpm until stationary phase. Subsequently cells were harvested and lysed and the genomic DNA was extracted using the DNeasy® Blood & Tissue Kit (Qiagen), according to manufacturer’s instructions. Plasmid DNA from both the ETEC strains was additionally isolated using the Gene Jet Plasmid Miniprep Kit (Thermo Scientific) by following the manufacturer’s instructions.

Genome sequences were assembled using SPAdes^84^ for *V. cholerae* V52 and SPades and plasmidSPAdes^85^ for ETEC E616 and ETEC E36. PATRIC^60^ and eggNOG mapper^86^ were used for genome annotation.

### Reconstruction of *V. cholerae* GEM, iAM-Vc960

A list of metabolic pathways in *V. cholerae* V52 was built based on the genome annotation generated in this study as well as those available in PATRIC and that of *V. cholerae* O1 N16961 (see Table S1at https://github.com/alyamahmoud/coinfection_modeling/blob/master/supplementary_material/supplementary_tables.xlsx). The reconstruction was converted into a model and the stoichiometric matrix was constructed with mass and charge balanced reactions in the standard fashion using the COBRA toolbox v.3.0.^36^. Flux balance analysis was used to assess network characteristics and perform simulations^37^. We used *i*JO1366^38^ as a starting point for reconstruction efforts which is a common practice to use the closest available species as a starting template^13,14^ while keeping only reactions for which evidence exists to be present in *V. cholerae* genome and/or transcriptome (see Table S1 at https://github.com/alyamahmoud/coinfection_modeling/blob/master/supplementary_material/supplementary_tables.xlsx). We also built an objective biomass function based on *i*JO1366 and *V. vulnificus*^7^ previously reconstructed GEMs. Additional reaction content was added from KEGG, and BIOCYC databases. All reactions added were manually curated according to a published protocol^83^. *i*AM-Vc960 was assessed for mass balance^83^. Metabolites charges and formulae were obtained from BiGG^87^ and updated in *i*AM-Vc960 to mass-balance the respective reactions. All reconstruction, refinement, validation and simulations using all models in this study were done using the COBRA toolbox^36^ (v3.0.) and Matlab-R2016b. Please refer to section “Refinement of *i*AM-Vc960” in the supplementary text at https://github.com/alyamahmoud/coinfection_modeling/blob/master/supplementary_material/supplementary_text.docx for more details on the curation steps of *i*AM-Vc960.

### Validation of *i*AM-Vc960 single gene deletion essentiality predictions

We downloaded gene essentiality data for *V. cholerae O1* str. C6706 from the Online GEne Essentiality (OGEE) database^4,5^. In total, 3886 genes (total number of ORFs identified in *V. cholerae*) were tested for essentiality. 458 genes were essential, 148 were essential for fitness, 3144 were non-essential and 136 were unknown. Out of the 458 essential genes, 145 were metabolic genes and were already in *i*AM-Vc960. *i*AM-Vc960 predicted 94 of those to be essential while the remaining 51 were falsely predicted by the model as non-essential. For the non-essential genes, 758 of those were already in *i*AM-Vc960. The model could predict 693 as non-essential while 65 were falsely predicted by the model as essential. The overall accuracy of the model predicted single gene essentiality was 87% (Figure 2C). This discrepancy between the model predictions and the OGEE dataset, the high confidence set that we used earlier and assuming low experimental error rate, indicates that the reconstructed *V. cholerae* reactome is incomplete and that there is further room for improvement and refinement of the *i*AM-Vc960 representing opportunities for new biological discoveries.

### Metabolic modeling of co-infection of *V. cholerae* and ETEC

To simulate co-infection, individual species models were combined into a community model where each species would interact with a common external metabolic environment through their metabolite exchange reactions^58,59^. This allowed each species to access the pool of media/infection site metabolites as well as metabolites that are released/uptaken by the other pathogen. Each species could secrete/uptake only those metabolites for which an exchange reaction (e.g. via transporters or free diffusion) exists in the model. The widely-employed FBA objective of biomass maximization^37^ was replaced with the maximization of a weighted sum of the biomass production fluxes for the community members^88^, i.e. the objective function was set to maximize the biomass function of each pathogen, simulating growth at 1:1 species composition/abundance. Flux balance analysis (FBA) was performed using open CORBA in Matlab 2016b, and the Gurobi solver v7.0. Please refer to section “Quality control of the Co-culture model *i*Co-Culture2993” in the supplementary text for more details on the curation of the co-culture model.

### Catabolic capabilities of *V. cholerae, ETEC* and co-infection GEMs

Growth in 656 different growth supporting conditions was simulated for *i*AM-Vc960, *i*ETEC1333 and *i*Co-Culture and then compared to identical simulation conditions for 55 GEMs of *E. coli* and *Shigella*^13^. Table S4 at https://github.com/alyamahmoud/coinfection_modeling/blob/master/supplementary_material/supplementary_tables.xlsx details the simulation conditions for the alternative nutrient sources and Table S5 at https://github.com/alyamahmoud/coinfection_modeling/blob/master/supplementary_material/supplementary_tables.xlsx shows all simulated growth conditions. Nutrient sources with growth rates above 0.01 were classified as growth supporting, whereas nutrient sources with growth rates less than 0.01 were classified as non-growth supporting. The binary results from the growth/no growth simulations were used to reconstruct the heatmap (Figure 3C). Ward’s agglomerative clustering of the matrix of correlations was used to cluster the species. The heatmap was visualized using the pheatmap R package. The ternary plot (Figure 4B) was visualized using the ggtern R package^89^.

### RNA Extraction, Sequencing and Data Analysis

Sampling of cells for the purpose of RNA extraction was performed as follows: Bacterial cells (mono-cultures and co-cultures of *V. cholerae* and ETEC) were grown to mid logarithmic phase in shake flasks at 37 °C at 200 rpm. In case of the co-cultures, equal concentrations of individual mono-cultures were inoculated into the same medium from the start. Once the appropriate growth phase was reached, the cells were harvested. RNA was extracted from the harvested cells using the RNeasy^®^Mini Kit (Qiagen), according to manufacturer’s instructions. Experiments were carried out in triplicates. The RNA extracted was in the range of 200 – 100 ng/µL.

RNAseq reads from mono-cultures were directly aligned to the genome assembly of the corresponding species. To check for reads cross mapping, we first attempted to map *V. cholerae* reads against ETEC genome assembly and vice versa. In either case, the percentage of mapped reads was < 2% (see Figure S4 at https://github.com/alyamahmoud/coinfection_modeling/blob/master/supplementary_material/supplementary_text.docx) indicating minimal cross-mapping between the two species. Following, we constructed an artificial genome assembly of both *V. cholerae* and ETEC combined, i.e. representing the co-culture as a single entity by merging the genome assemblies of the two species. PATRIC^60^ was used for annotation of the merged genome assembly. *V. cholerae* and ETEC reads from the co-culture were then each separately aligned against the merged genome assembly and read counts were computed, i.e. we sequenced and annotated the genome sequences from the single and dual cultures using the same assembly and annotation pipeline to avoid differential gene calling. Although all strains used in this study (*V. cholerae* V52, ETEC E36 & E616) are clinical isolates that have been sequenced and characterized before^62,63^, we have generated new assemblies and annotations mainly for the sake of consistency for gene calling where we subjected the mono- and co-culture transcriptomes to the same processing and annotation pipelines. Bowtie2^90^ was used for all genome alignment. Read counts for all genes were extracted with HTSeq-count^91^, normalized and analysed using the R package DESeq2^92^. In order to do differential expression analysis between the genome assemblies generated from the mono cultures and the co-cultures, we aggregated genes by their FIGfam IDs^93^. Members of a FIGfam, are believed to implement the same function, they are believed to derive from a common ancestor, and they can be globally aligned. We wanted to see if there are specific functions that will be significantly altered between the two culture conditions especially that the sequence identity between ETEC and *V. cholerae* is around 80%^43^. FIGfam IDs were aggregated by keeping the FIGfam ID with the maximum value of raw read counts across all replicates from both the mono- and co-cultures. GOstats^94^ R package was used for the GO enrichment analysis and GOplot^95^ R package was used for visualization of GO enrichment results in Figure 5. The details of procedure for dual RNAseq data analysis are outlined in Figure S4 and in the supplementary text at https://github.com/alyamahmoud/coinfection_modeling/blob/master/supplementary_material/supplementary_text.docx and code at the github repository at https://github.com/alyamahmoud/coinfection_modeling.

### Data availability

All data generated in this study are included in this published article. Models, supplementary test and supplementary tables as well as code to reproduce the main figures and key analyses in this study are available as part of a github repository at https://github.com/alyamahmoud/coinfection_modeling.

## Acknowledgements

We are very grateful to insightful comments from Dr Nathan Lewis and Dr Neema Jamshidi. We thank Dr Abdallah Abdallah and Mohammed Alarawi from the bioscience core lab at KAUST, Dr Hajime Ohyanagi and Dr Yoshi Saito for helpful discussions. This research was supported by funding from KAUST: BAS/1/1624-01-01, BAS/1/1059-01-01 and SEED funding: FCS/1/2448-01-01 (AM, XG, TG, and KM) as well as by grants from the Novo Nordisk Foundation (IM, VR), the Swedish national research council (VR) and the Danish national research council (DFF) (IM).

## Author contributions

AM performed the modeling, simulations, data analysis, and wrote the paper. VR performed the experiments. BJ provided support for the modeling and data analysis. JN provided support for the modeling. KM provided support for the DNA and RNA sequencing. XG contributed to data analysis. IM and TG conceived the project, oversaw the project and wrote the paper. All authors read and approved the final manuscript.

